# Smaller stepping thresholds in older adults might be related to reduced ability to suppress conflicting sensory information

**DOI:** 10.1101/2025.06.25.661273

**Authors:** Hannah D. Carey, Friedl De Groote

## Abstract

Aging leads to alterations in the sensorimotor system and balance control but it is not well understood how changes in sensorimotor function affect how people respond to postural disturbances. Elucidating the relationships between balance control and sensorimotor function is crucial for developing effective rehabilitations. Here, we compared the kinematic responses to platform translations and rotations during standing in 10 young and 30 older adults and explored relationships between sensorimotor function and balance responses. We found that older adults were less able to withstand perturbations without stepping, not because their non-stepping strategies were less effective but because they chose to step at smaller deviations of the extrapolated center of mass. Older adults performed worse than young adults on measures of sensory and motor function but lower stepping thresholds were associated with susceptibility to unreliable visual information and not with reduced sensory acuity or reduced strength. Poor sensory reweighting may contribute to and combine with age-related cognitive rigidity, leading to a higher priority on safer strategies. Older adults may resort to stepping, even if a step is not necessary, rather than rely on potentially inaccurate sensory signals to inform a corrective response. Our results provide initial evidence that sensory reweighting could be a potential target for fall prevention methods.

**NEW & NOTEWORTHY:** The relationship between age-related changes in sensorimotor function and postural control is poorly understood. Here, we did a comprehensive assessment of sensorimotor function and reactive standing balance. We found that healthy older adults chose safer strategies, i.e. they step at smaller disturbances, than young adults. Although we found many differences in sensorimotor function, only a reduced ability to suppress conflicting sensory information was related to the use of a safer strategy.

## INTRODUCTION

Potentially destabilizing conditions, such as a jostling bus, cracked pavement, or dim lighting, are common in daily life. Healthy people can typically withstand most of these disturbances, but changes in sensorimotor function that occur with normal aging leave older adults more vulnerable to falls. Aging deteriorates the sensory systems required to detect disturbances, the central processing used to interpret information and plan responses, and the motor capabilities needed to respond effectively(1). The process of sensory integration for postural control(2,3) and the impacts of aging on sensory systems(4–6) and balance(7–10) have been characterized but it is not well understood how changes in sensorimotor function affect how people respond to postural disturbances. This impedes both fall prevention and rehabilitation efforts, making it difficult to identify those at risk of falling and effectively target the underlying changes in balance control that may increase fall risk(11,12). Elucidating the relationships between balance control and sensorimotor function is thus crucial for understanding how normal aging leads to falls and developing strategies to mitigate this risk.

Though the specific mechanisms are not yet understood, the impacts of age-related sensorimotor changes can be seen in older adults’ balance control. Balance-correcting responses typically incorporate three main types of kinematic strategies, which have been well described(13,14). Minor disturbances may be avoided by using small movements at the ankle to adjust the center of pressure, referred to as the *ankle* or *center-of-pressure strategy*. Challenging disturbances may elicit more movement in the trunk, usually referred to as a *hip strategy*. If the ankle and hip strategies are insufficient, taking a step instantly widens the base of support to prevent a fall. Older adults are more likely to resort to large corrective responses like stepping in response to a disturbance(8,15), which has been attributed to age-related changes in both sensory noise(15) and muscle strength(16).

Older adults may step more often because sensory noise and delays interfere with accurate detection, muscle strength is insufficient to produce an effective response, or a safer strategy is preferred even if they are physically capable of withstanding the perturbation with a non-stepping strategy due to concerns about the consequences of a fall. Aging produces noise and delays in sensory signals(17,18) and degrades the individual proprioception, vision, and vestibular systems used to monitor posture(1). For example, age-related changes in muscle spindles and nerve transmission reduce the quality of proprioceptive information(4), leading to greater reliance on vestibular(1) and visual information(19), which are less sensitive to postural changes(20) and susceptible to aging themselves(1). Though people can readjust based on sensory information available (i.e., sensory reweighting(2)), reweighting is slower in older adults(21,22). Because age degrades multiple aspects of sensorimotor function simultaneously, and people often adapt to those changes, it is difficult to untangle the direct impacts of each change on balance control.

The characteristics of the perturbation will shape the sensory inputs and might thus influence the sensitivity to age-related changes in sensory acuity. In addition to proprioception, vestibular and visual information likely play important roles in shaping the response to support-surface translations and rotations that cause large head and trunk movements(23,24). Detecting small postural disturbances relies mainly on proprioception given its high acuity but the Small support-surface rotations primarily displace the ankle, with little impact on trunk orientation(23); detecting small rotations is likely most dependent on proprioception. Because trunk orientation can be estimated from visual, vestibular, and proprioceptive systems provide additional information when larger disturbances occur. When standing is perturbed by support surface translations, changes in body orientations can be derived from proprioceptive information alone but additional information from the visual and vestibular systems increases accuracy. In contrast, proprioceptive information alone is not sufficient to infer information about trunk orientation during support surface rotations. Responses to rotational perturbations may therefore be less robust against decline in visual and vestibular function. We can therefore use different types of perturbations to probe the role of sensory function in balance control.

Accordingly, our objectives for this work were to compare the kinematic responses to platform translations and rotations in young and older adults and to look for relationships between sensorimotor function and balance responses. We hypothesized that older adults would step more often, exhibit larger trunk and ankle movements, and score lower in sensory and motor function measures than young adults, as in previous literature. Because we did not have supporting evidence for which aspects of sensorimotor function might be important for balance control, our analysis of associations between sensorimotor function and balance control was exploratory.

## MATERIALS AND METHODS

### Participants

All experimental protocols were approved by the ethical committee of UZ/KU Leuven (Protocol No. 61361). Ten healthy young adults and thirty healthy older adults (Table 1, Figure 1 -A) provided written informed consent before participating in this study.

**Table 1.**
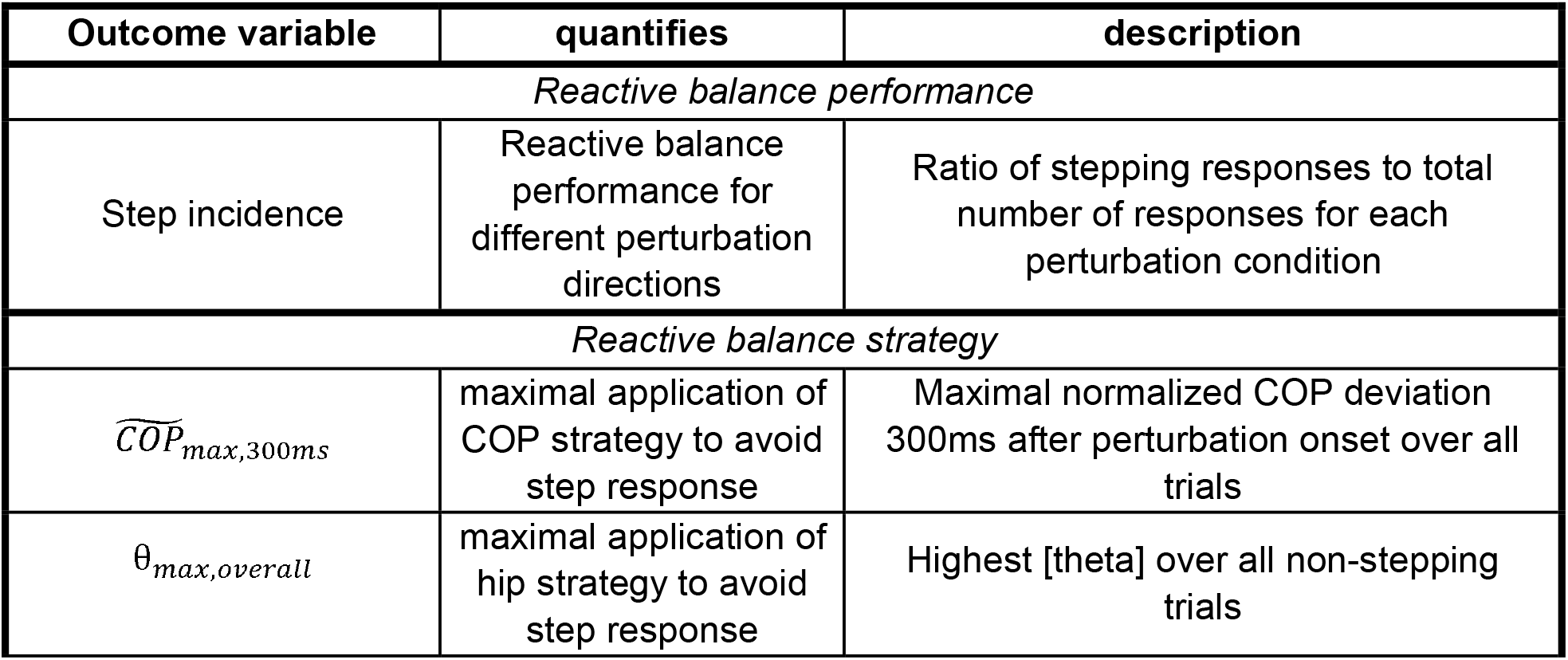
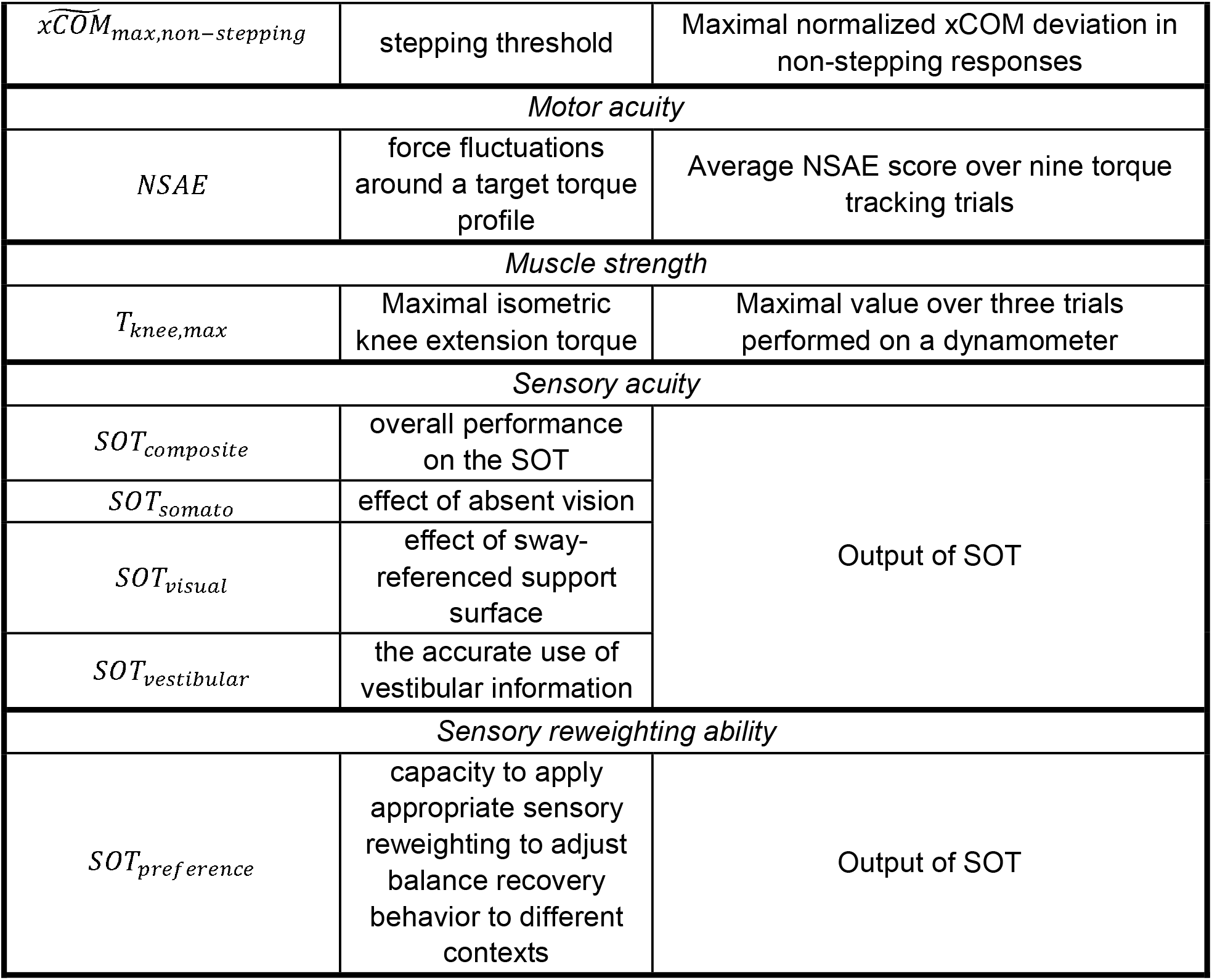
Effect of ANG II infusion on biomechanical properties of parenchymal arterioles.

**Figure 1.**
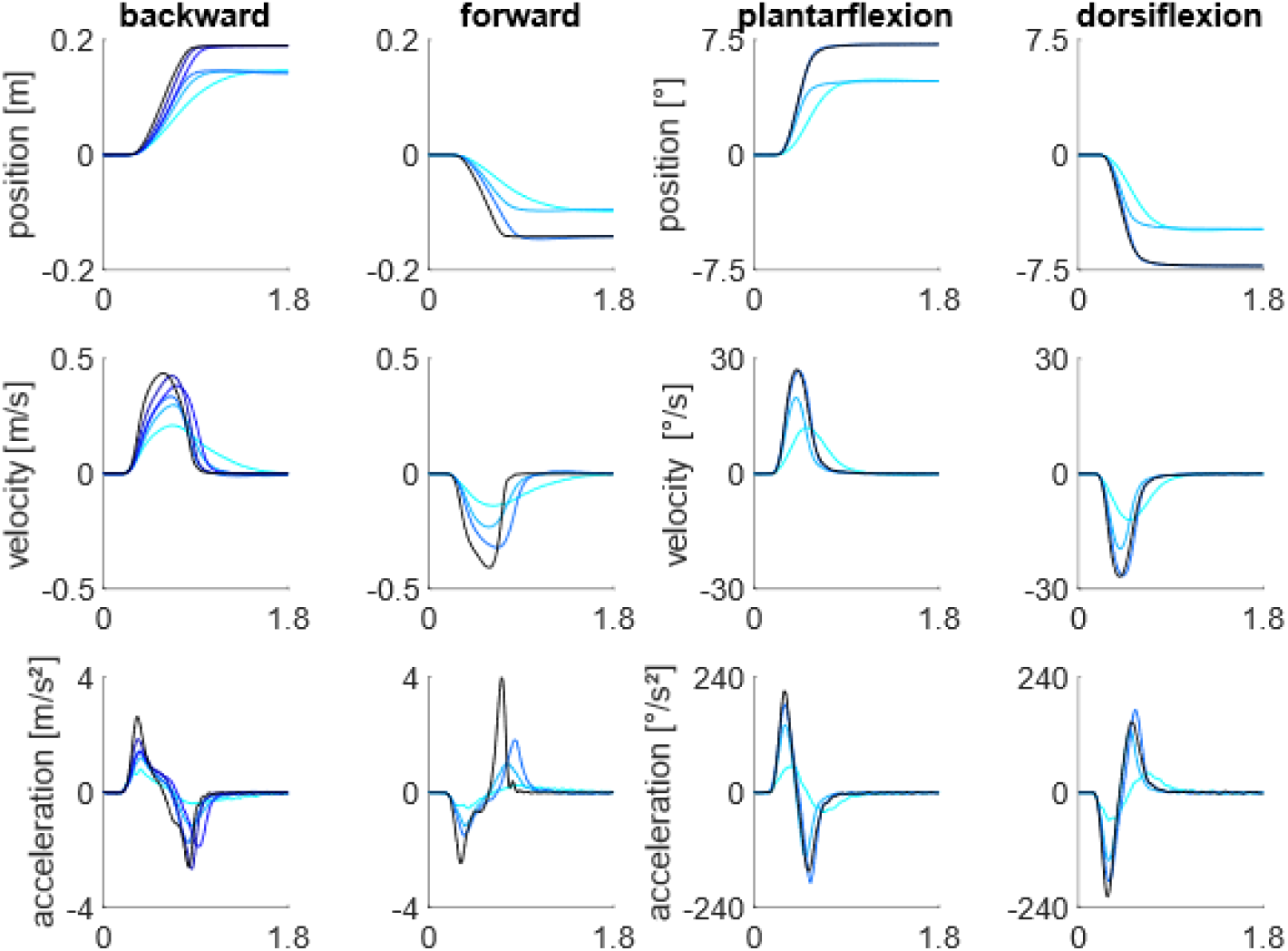
Position, velocity, and acceleration profiles for the backward support-surface translations, forward support surface translations, plantarflexion pitch rotations, and dorsiflexion pitch rotations

Participants had not been enrolled in similar reactive balance assessments previously. Participants who suffered from musculoskeletal injury or pathologies that could impair balance were excluded. To qualify the group of older adults as healthy, they performed a 5x sit-to-stand test(25) and a test measuring handgrip strength(26) and filled out the Fall Efficacy Scale-International (FES-I) questionnaire(27,28). None of the included older adults scored under the cut-off points for the 5x sit-to-stand test or hand grip strength test that would indicate frailty(29). None of the participants had a ‘high concern’ score for the FES-I.

We quantified standing reactive balance performance and kinematic strategies in response to perturbations applied by platform translations and rotations (CAREN platform, Motek), muscle strength (Biodex), motor acuity (Biodex), sensory acuity (NeuroCom Balance Master), and sensory reweighting capacity (NeuroCom Balance Master). Table 2 provides a summary of the outcome variables.

### Standing Reactive Balance

#### Protocol & data processing

Surface perturbations were applied to participants during standing to assess reactive balance(30). Perturbations were applied as support-surface translations and rotations in different directions using a CAREN platform (Motek Medical, Netherlands). Participants stood barefoot on the movable platform with their feet at shoulder width, looked forward, and wore a safety harness to catch them in case of a loss of balance. Participants were instructed to maintain balance without taking a step when perturbed and were allowed to move their arms freely. If the perturbation elicited a stepping response, participants were instructed to return to their original position before the next perturbation. To standardize foot placement, the heel position was marked on the platform. Participants received support-surface perturbations in six directions and with different magnitudes (Figure 1): anterior and posterior translations, lateral left and right translations, and pitch rotations in two directions, inducing ankle plantar-or dorsiflexion. Lateral translations were included to prevent subjects from anticipating the perturbation directions; these trials were not included in any analyses. The protocol consisted of a familiarization part and a randomized part. Subjects were first familiarized with the platform motion while being informed on the direction of the upcoming perturbation. During the familiarization, perturbations were applied with progressively larger magnitudes until subjects needed to take a step, which ended familiarization with the specific perturbation direction. The first perturbation magnitude that induced a step response was the highest magnitude included in the second randomized part of the protocol. When no step response was evoked at the highest magnitude, all perturbations for that direction were included. Up to six different perturbation magnitudes were presented for posterior translations, whereas up to four different perturbation magnitudes were presented in the other directions (Figure 1). Next, during the randomized part of the protocol, each perturbation condition was applied five times in random order. Perturbations were provided at random to minimize anticipatory postural adjustments. Motion capture data was collected: subjects were instrumented with 33 reflective markers on anatomical landmarks (full body plug-in-gait) and cluster markers on the left and right shanks and thighs. Platform motion during perturbations of standing was measured using three markers. The marker trajectories were captured using seven Vicon cameras at a frequency of 100Hz. The CAREN platform was instrumented with two force plates, measuring contact forces and moments between the subjects and the support surface at 1,000Hz. A static trial in anatomical position was acquired before starting the experiments.

Data was preprocessed to get joint, center of mass (COM), and center of pressure (COP) kinematics and joint kinetics. All marker trajectories were labeled in Vicon Nexus 2.4. Generic musculoskeletal models (gait2392 - OpenSim 3.3) were scaled based on the subject mass and anatomic marker positions acquired during the static trial(31,32). Joint angles were computed using OpenSim’s Inverse Kinematics tool (OpenSim3.3). Finally, OpenSim’s Body Kinematics tool was used to compute segment and whole-body kinematics. COP trajectories were derived from the forces and moments recorded by the force plates. Joint kinetics were computed using OpenSim’s Inverse Dynamics tool (OpenSim3.3), with the scaled musculoskeletal models, force plate data, and joint kinematics as input. A correction of the force plate data was performed to remove forces and moments registered due to the inertia of the force plate(33). We corrected for these forces and moments by subtracting the forces and moments registered while the platform was moving without any load from the data acquired with the subject on the platform(30).

Perturbation onsets were identified based on the acceleration of a marker fixed to the treadmill. For each trial, we identified “trial onset” as the first frame where the acceleration of the marker was more than three standard deviations larger than during a baseline window prior to the perturbation. This trial onset was manually adjusted if identification was poor (5 trials total). We then calculated the median of trial onsets within each perturbation type and used that value as the new perturbation onset for all trials. This method produced better alignment of platform and participant motion profiles than using individual onsets for each trial.

### Step incidence to quantify reactive balance performance

During perturbed standing, step incidence within each perturbation type (direction and magnitude) was computed by detecting trials where the vertical ground reaction force was below 10N for more than 50ms.

### Outcome variables to quantify reliance on balance-correcting strategies

To quantify the reliance on COP strategies, we evaluated the center of pressure shift during the automatic response (, with representing the COM height during quiet standing). We defined the use of a COP strategy as the maximal magnitude of anteroposterior center of pressure shift achieved during the automatic response following platform perturbations over all trials, with maximum anterior position used for backwards translations and toe-down rotations, and maximum posterior position used for forwards and toe-up rotations.

To quantify the reliance on hip strategies, we computed the overall highest value of the maximal trunk lean angle in the sagittal plane () across all non-stepping responses for each perturbation direction.

The reliance on a stepping strategy was quantified by the stepping threshold, which was defined as the maximal extrapolated center of mass excursion in non-stepping responses. This outcome variable captures how strongly a subject’s balance was disturbed before they initiated a stepping response. For each non-stepping trial, we computed the largest within trial extrapolated center of mass excursion, xCOM. For each subject, we computed the mean over

the three largest values *xCOM*_*max*,*non-stepping*_. We chose to take the mean over the three largest values to avoid accidental outliers not representative of the overall subject behavior. We tested whether taking the average over the largest four or five values influenced our conclusions, which was not the case. xCOM positions were normalized by COM height during quiet standing(34): 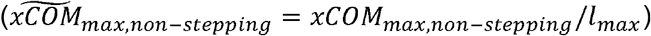.

### Measures of sensory and motor function

#### Muscle strength

Muscle strength was assessed for the knee-extensor muscles on a Biodex System 4 PRO dynamometer (Shirley, NY). Subjects were seated on the Biodex equipment with their knee at a 90° flexion angle with the back support in fully upright position. Subjects performed three maximal voluntary isometric knee extension contractions of three seconds with one-minute rest between trials. The torque was measured at 1000Hz. From these three trials the maximal voluntary isometric knee extension torque, normalized by the body weight, was computed (*T*_*knee*,*max*_).

#### Motor acuity

Motor acuity was tested by measuring force fluctuations during submaximal isometric knee extension(18,35,36). Force fluctuations were tested at 15% and 20% of the maximal isometric knee extension torque measured. To assess force fluctuations, three different torque tracking tasks were executed in random order. Task 1 and 2 consisted of generating a constant torque for 15s at respectively the 15 and 20% level, task 3 consisted of tracking a ramp-up torque from the 15 to 20% level during 15s. The target torque profile was displayed on a monitor and participants were instructed to match the torque level as well as they could for the duration of each test by generating knee-extension torque at 90° flexion angle. The torque generated by the subjects (GT) was overlaid in real time on the target torque.

The recorded torque profiles were low-pass filtered using a fourth-order low-pass Butterworth filter with a cut-off frequency of 25Hz(35). Force fluctuation for each of the trials was computed using the normalized standard deviation (SD) of the absolute error (NSAE) between the target and generated torque during the middle ten seconds of each trial(36,37):

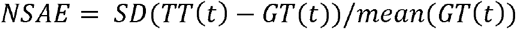

A composite score to quantify motor acuity for each subject was then computed by taking the average over the nine trials.

### Sensory acuity & sensory reweighting capacity

Sensory acuity of the visual, vestibular and proprioceptive systems as well as the capacity to perform sensory reweighting was assessed using a sensory organization test (SOT)(7,38–40). During SOT participants are instructed to stand as still as possible in six different conditions. Visual and somatosensory stimuli are manipulated throughout the six conditions of the test, each of which consists of three 20-second trials. In condition 1, the control condition, all sensory information is available. The participant looks straight ahead. In condition 2, participants close their eyes, eliminating visual input. In condition 3, the visual surround is sway-referenced such that the visual information becomes unreliable and in conflict with vestibular and somatosensory information. Conditions 4, 5 and 6 repeat conditions 1, 2 and 3 but now the support-surface rotates in phase with the subject’s postural sway, making somatosensory information, mainly from the ankle joint, unreliable and in conflict with other sensory information.

During the SOT, anteroposterior sway of the body is computed based on the measured center-of-pressure movement and an approximation of the human body as a single inverted pendulum. Both visual and support-surface sway-referencing is done based on the estimated anteroposterior sway angle (*θ*_*AP*_). For each trial an equilibrium score is computed based on the amplitude (*θ*_*AP*,*max*_ *-θ*_*AP*,*min*_) of the estimated anteroposterior sway. A linear scoring system is applied where an anteroposterior sway amplitude of 12.5° leads to an equilibrium score of 0 and an AP sway amplitude of 0° generates an equilibrium score of 100. Trials with anteroposterior sway amplitudes that exceed 12.5° or failed trials where the feet were not kept in place generated an equilibrium score of 0. The average equilibrium score (EQ) over the three trials is computed for each condition. Five SOT scores are then computed from these conditions specific equilibrium scores:

- **Composite score** (*SOT*_*composite*_), a measure for overall performance on the test.
- **Somatosensory score** 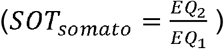, a measure of the effect of absent vision, which may reflect accurate use of somatosensory information.
- **Visual score** 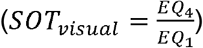, a measure of the effect of unreliable somatosensory information, which may reflect accuracy of visual information.
- **Vestibular score** 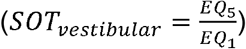, a measure of the effect when only vestibular information is reliable (absent vision and unreliable somatosensory information).
- **Preference score** 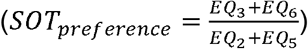, a measure of the ability of central adaptive mechanisms to suppress conflicting visual input, quantifies the capacity to apply appropriate sensory reweighting to adjust balance recovery behavior to different contexts.

### Statistics

The sample size was chosen to at least have 80% power at the 0.05 alpha level when differentiating older and young adults based on strength, sensory, and motor acuity, measures that were found to have large effect sizes (Cohen’s d = 0.9) in previous studies(41). No a-priori power analysis was conducted for the correlation analysis. Comparisons of stepping incidence between young and older adults were performed in jamovi (The jamovi project); all other statistics were performed in MATLAB (Mathworks).

### Comparison of stepping incidence between young and older adults

We compared step incidence across perturbation directions, difficulty levels, and age groups using a mixed three-way ANOVA. Because few older adults could perform the largest difficulty level in each direction, these levels were excluded. For the backwards perturbations, we used the first (smallest), third, and fifth difficulty levels, to maintain a balanced ANOVA design. Post hoc tests were performed with Bonferroni corrections.

### Comparison of sensorimotor function and balance-correcting response between young and older adults

We applied parametric and non-parametric, depending on normality of the data which was tested using a Shapiro-Wilkes test, one-tailed unpaired t-tests to test whether *SOT*_*composite*_, *SOT*_*somato*_, *SOT*_*visual*_, *SOT*_*vestibular*_, *SOT*_*preference*_, *T*_*knee*,*max*_, and *NSAE* were significantly larger in older adults than in young adults. We followed the same approach to compare stepping thresholds, ankle response, and hip response between young and older adults. To account for multiplicity, we applied the two-step method that controls the false discovery rate when testing multiple hypotheses from Benjamini et al.(42,43). This method is less conservative than Bonferroni corrections that over correct when tests might be positively correlated. Differences between young and older adults with large effect sizes have been found in previous studies for these factors(35,38,44). As such our sample size sufficed to have 80% power at the 0.05 α level assuming a large effect size (Cohen’s d = 0.9).

### Exploratory assessment of associations between measures of sensorimotor function and balance-correcting response

We investigated whether any measures of sensorimotor function were associated with either stepping threshold, ankle, or hip response. Before performing the linear regressions, outliers in sensorimotor function and balance variables were removed. Outliers were defined as values larger than the 75^th^ percentile + 1.5 interquartile ranges, or lower than the 25^th^ percentile - 1.5 interquartile ranges. For each balance and sensorimotor function variable, we removed up to 5 outlier subjects. We formulated linear models as *b*_*var*_ *∼ s*_*var*_ *+ s*_*var*_ *· age group*, where *b*_*var*_ indicates the balance response variable (in a given perturbation direction), *s*_*var*_ represents the sensorimotor function variable, and *age group* is a binary variable representing the age groups. We used a separate linear model for each sensorimotor function, balance response variable, and perturbation direction. For the comparisons in which the whole model was significant (p<0.05), we used an ANOVA to evaluate the contribution of each model term. As we were interested in the influence of sensorimotor function on balance responses, we dismissed models in which age group was the only significant factor and visual inspection revealed two clouds of data suggesting that both balance responses and sensorimotor function changed with age but were not otherwise related.

## RESULTS

### Older adults step more often than young adults during larger perturbations

Older adults stepped more often than young adults during backward, forward, and dorsiflexion perturbations, but only at higher perturbation magnitudes. Few participants from either group stepped during the low magnitude perturbations or the plantarflexion perturbations at any level.

A step occurred in only 11 plantarflexion trials across all subjects regardless of age, representing 2% of all plantarflexion trials. In contrast, steps occurred in 22% of all backward trials, 29% of all forward trials, and 11% of dorsiflexion trials. The same young adult stepped during one trial of the largest dorsiflexion and the largest plantarflexion perturbation; older adults made all other steps during rotational perturbations.

We identified significant interactions between difficulty and age group (F(2,72)=9.09), direction and age group (F(3,108)=4.12, p=0.008), and direction and difficulty (F(6,216)=18.08, p<0.001). There was no significant three-way interaction. Post hoc tests for perturbation direction indicated higher stepping incidence during backwards translations compared to dorsiflexion (p=0.001) and plantarflexion (p<0.001) rotations, and higher step incidence in forward translations compared to dorsiflexion (p<0.001) and plantarflexion (p<0.001) rotations (Figure 2, top). There was no significant difference in step incidence between backward and forward translations. Post hoc tests for perturbation difficulty level indicated that step incidence increase with perturbation difficulty. Step incidence was lower in small perturbations than medium (p=0.004) or large perturbations (p<0.001), and lower in medium than large perturbations (p<0.001) (Figure 2, bottom). See supplementary material for a detailed figure of step incidence (S1) and full results of the ANOVA and post hoc tests (S2).

**Figure 2.**
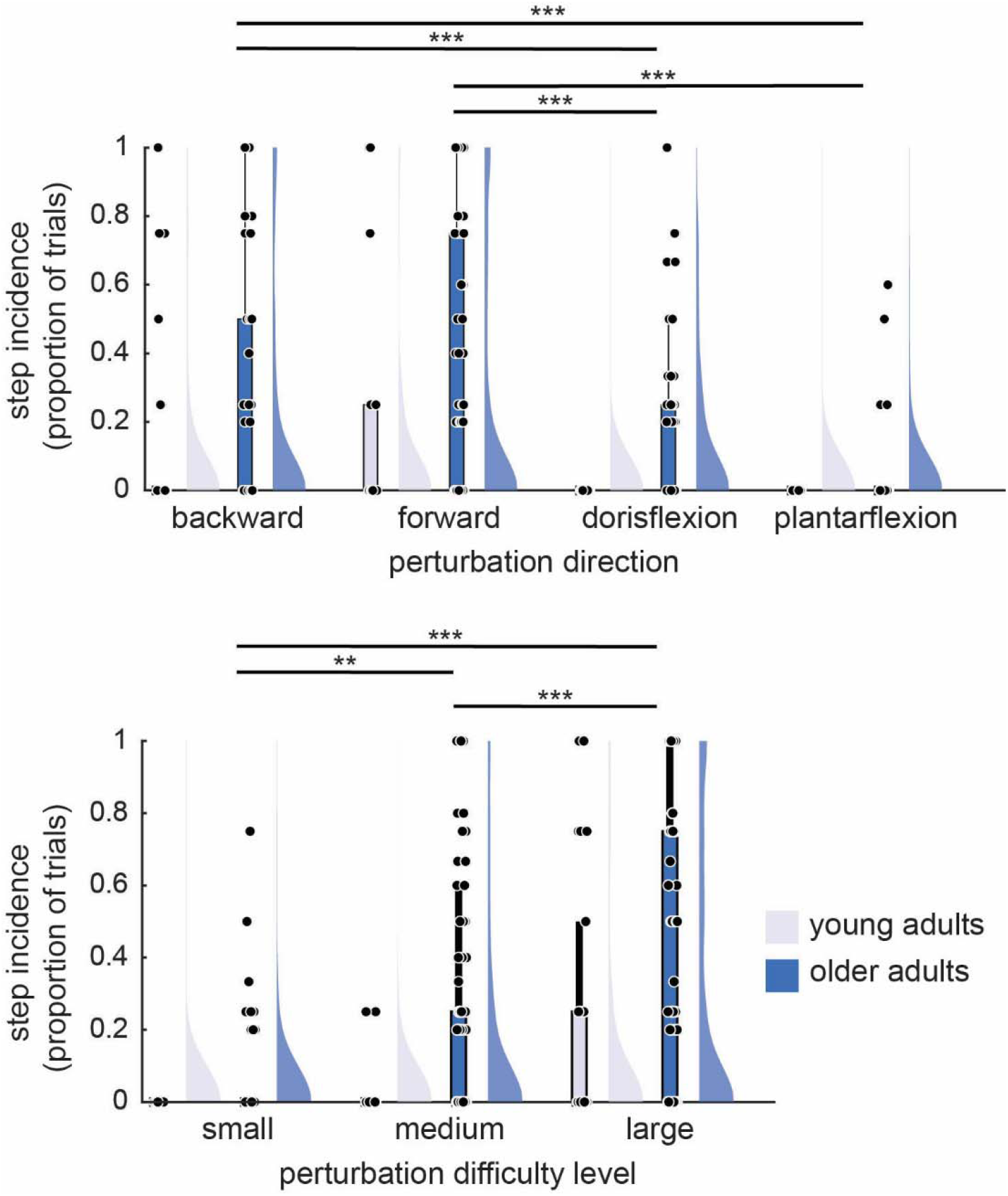
Raincloud plots of step incidence in young (light purple) and older adults (dark blue), separated by perturbation direction (top) and difficulty level (bottom). Dots represent the proportion of trials in which a participant stepped in each direction and difficulty level. Stars represent significance of post hoc tests for direction (top) and difficulty level (bottom), ** for p<0.01 and *** for p ≤0.001.

### Older adults had a smaller stepping threshold during translations, but higher thresholds during rotations

Older adults exhibited smaller stepping thresholds than young adults during backward (rank-sum, p=0.01) and forward (t-test, p=0.003) perturbations, as measured by the maximum xCOM deviation during non-stepping responses (Fig. 3, left). In contrast, older adults exhibited larger stepping thresholds during the rotational perturbations (dorsiflexion: t-test, p<0.001, plantarflexion: t-test, p<0.001, Fig. 3, left). Stepping thresholds during rotations were generally lower than stepping thresholds during translations.

**Figure 3.**
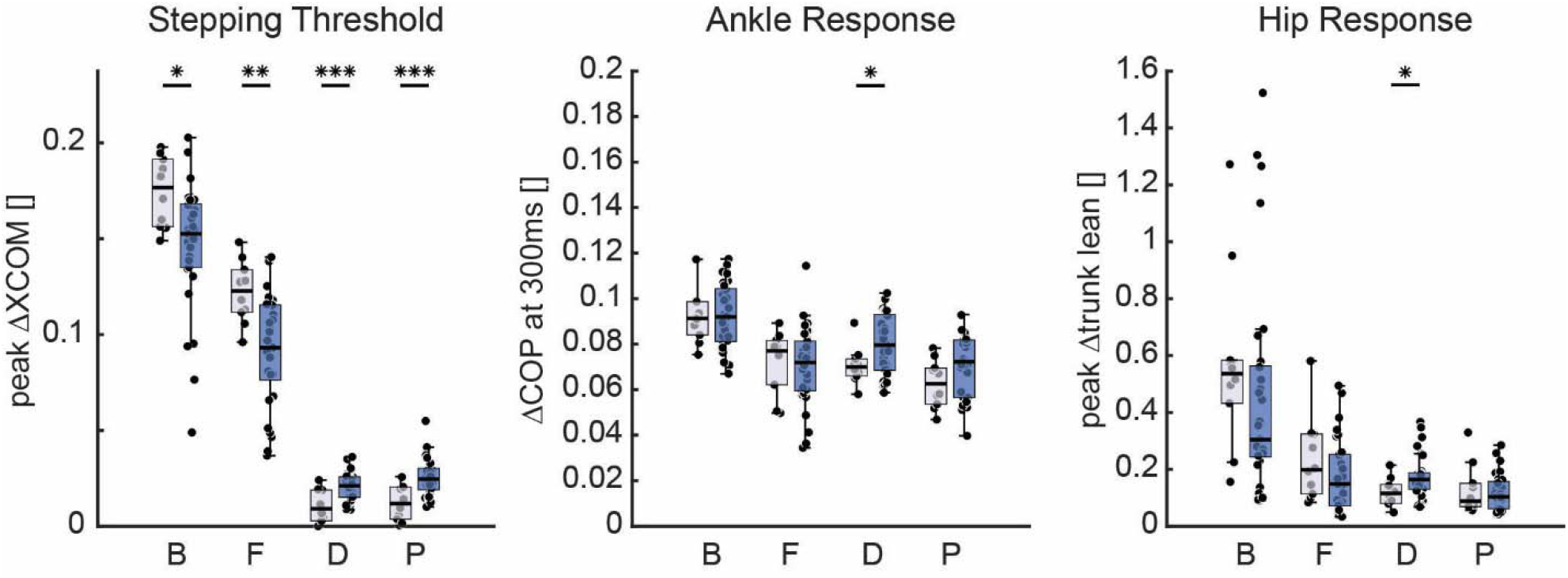
Measures of balance strategy in young and older adults: stepping threshold (peak magnitude XCOM during non-stepping trials, left panel), ankle response (change in anterior-posterior COP at 300 ms, middle panel), and hip response (peak magnitude of sagittal plane trunk angle in non-stepping trials, right panel), aggregated across difficulty levels for each perturbation direction. Box plots indicate the median and interquartile range, with young adults in light purple and older adults in dark blue. Statistically significant differences between young and older adults are indicated with stars. * for p<0.05, ** for p ≤0.01, and *** for p ≤0.001.

### Older adults use larger hip and ankle responses than young adults during dorsiflexion rotations

Older adults exhibited larger ankle responses in dorsiflexion perturbations than young adults (YA:0.071±0.008, OA:0.08±0.013, p=0.04). Young and older adults exhibited similar levels of ankle response during translations and plantarflexion perturbations (all t-tests, see table xx and Fig. 3, middle panel). Dorsiflexion perturbations also elicited larger hip responses in older adults than younger adults (rank-sum, median YA: 0.12, OA: 0.16, p=0.03). No significant differences in hip response were found in translation or plantarflexion perturbations (all rank-sum tests, see table I and Fig. 3, right panel).

### Older adults have decreased visual function, vestibular function, muscle strength and lower sensory reweighting capacities compared to young adults

As measured by the sensory organization test, older adults had decreased visual and vestibular function and lower sensory reweighting capacities compared to young adults. Although they were weaker, they had similar proprioceptive function and motor acuity to young adults.

Older adults had lower sensory organization test composite scores than young adults (YA: 82.8±4.16 vs OA: 70.0±7.94; p = 2.76e-5) (Figure 4 – COMP). The composite score was further differentiated into sub scores. First, older adults had significantly lower vestibular scores than young adults (YA: 76.4±7.41 vs OA: 54.7±15.4; p = 4.58e-5) (Fig. 4 - VEST), derived from a larger increase in body sway, with respect to the reference condition, when eyes closed and sway-referencing of the platform. Second, older adults had significantly lower visual scores than young adults (YA: 93.6±2.55 vs OA: 85.4±8.05; p = 9.83e-4) (Fig. 4 - VIS), indicating the sway-referenced support elicits greater postural sway in older adults than in younger adults. Third, older adults had significantly lower preference scores than young adults (YA: 100±6.12 vs OA: 92.3±11.8; p = 0.0299) (Fig. 4 - PREF), indicating a decreased capacity to suppress conflicting sensory input (i.e. sensory reweighting) in older adults. Finally, somatosensory scores were not significantly different between young and older adults (YA: 96.0±1.99 vs OA: 96.9±3.37; p = 0.318) (Fig. 4 - SOM).

**Figure 4.**
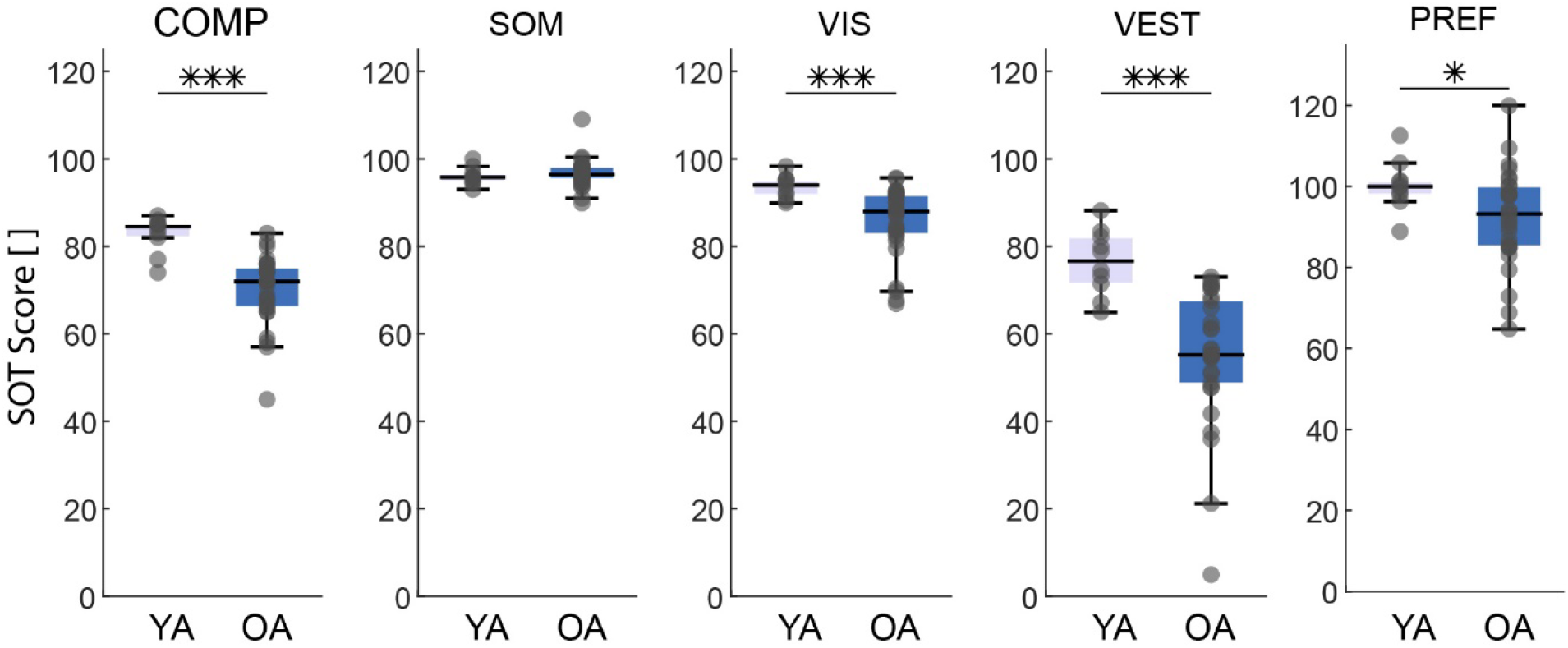
Sensory Organization Test (SOT) scores for the composite score (COMP) and the four subscores, in young (light purple) and older adults (dark blue). Stars represent significant difference between young and older adults, * for p<0.05 and *** for p ≤0.001.

Young adults were significantly stronger than older adults (Figure 5 - Left), but motor acuity did not differ between young and older adults. On average, young adults could generate 36% more normalized knee extension torque (YA: 3.42±1.31 Nm/kg vs OA: 2.19±0.57 Nm/kg; p = 1.90e-4). The normalized standard deviation of the absolute error (NSAE) for the isometric torque tracking task was not significantly different between young and older adults (YA: 0.0179±0.0036 vs OA: 0.0174±0.0041; p = 0.725) (Fig. 5 - right), indicating that peripheral motor acuity of the knee extensors at submaximal forces is similar between young and older adults.

**Figure 5.**
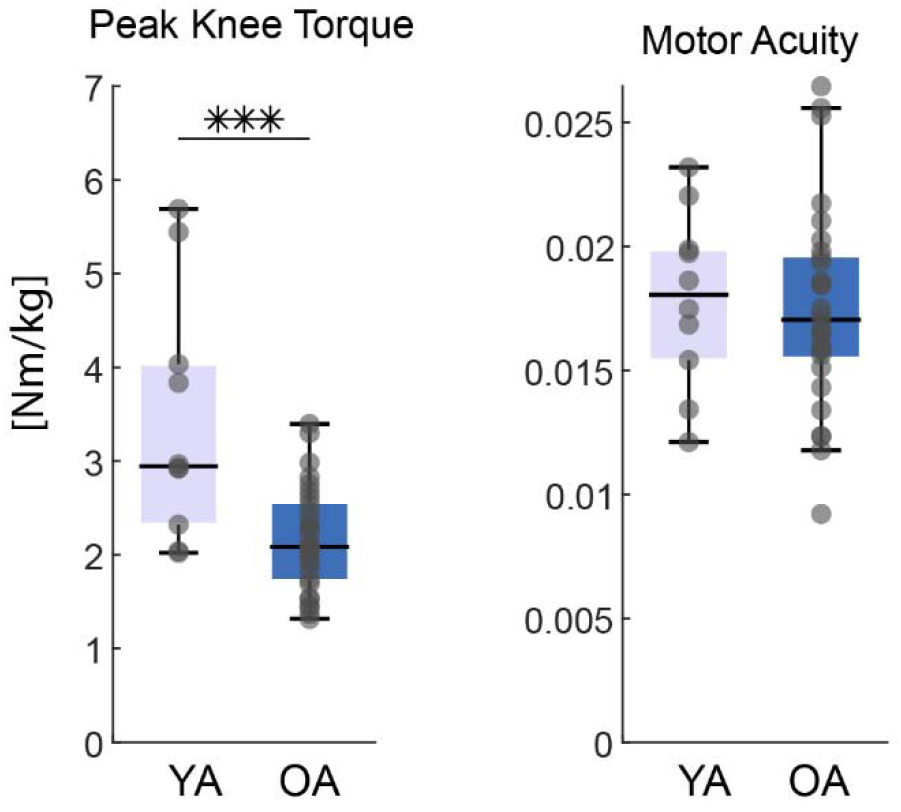
Measures of muscle strength, i.e. peak knee torque (left), and motor acuity, i.e. normalized standard deviation of absolute error (NSAE) of torque-matching task, (right) in young (light purple) and older adults (dark blue). Statistical significance indicated with stars, *** for p ≤0.001.

### Exploratory relationships between balance response and measures of sensorimotor function

We found seven significant relationships between balance responses and sensorimotor function. There was no significant interaction effect between age group and sensory function for three of these comparisons (shown as green blocks in Fig. 6). There, we reported the Pearson’s correlation coefficient between the balance and sensorimotor variables without regard to age group. For all comparisons with interaction effects, there was a relationship in young adults, but none in older adults. Accordingly, we reported the correlation coefficients for each age group. See Figure 6 (center) for a visual overview of the significant relationships. *More detailed results from this exploratory analysis can be found in the Supplementary Data*.

**Figure 6.**
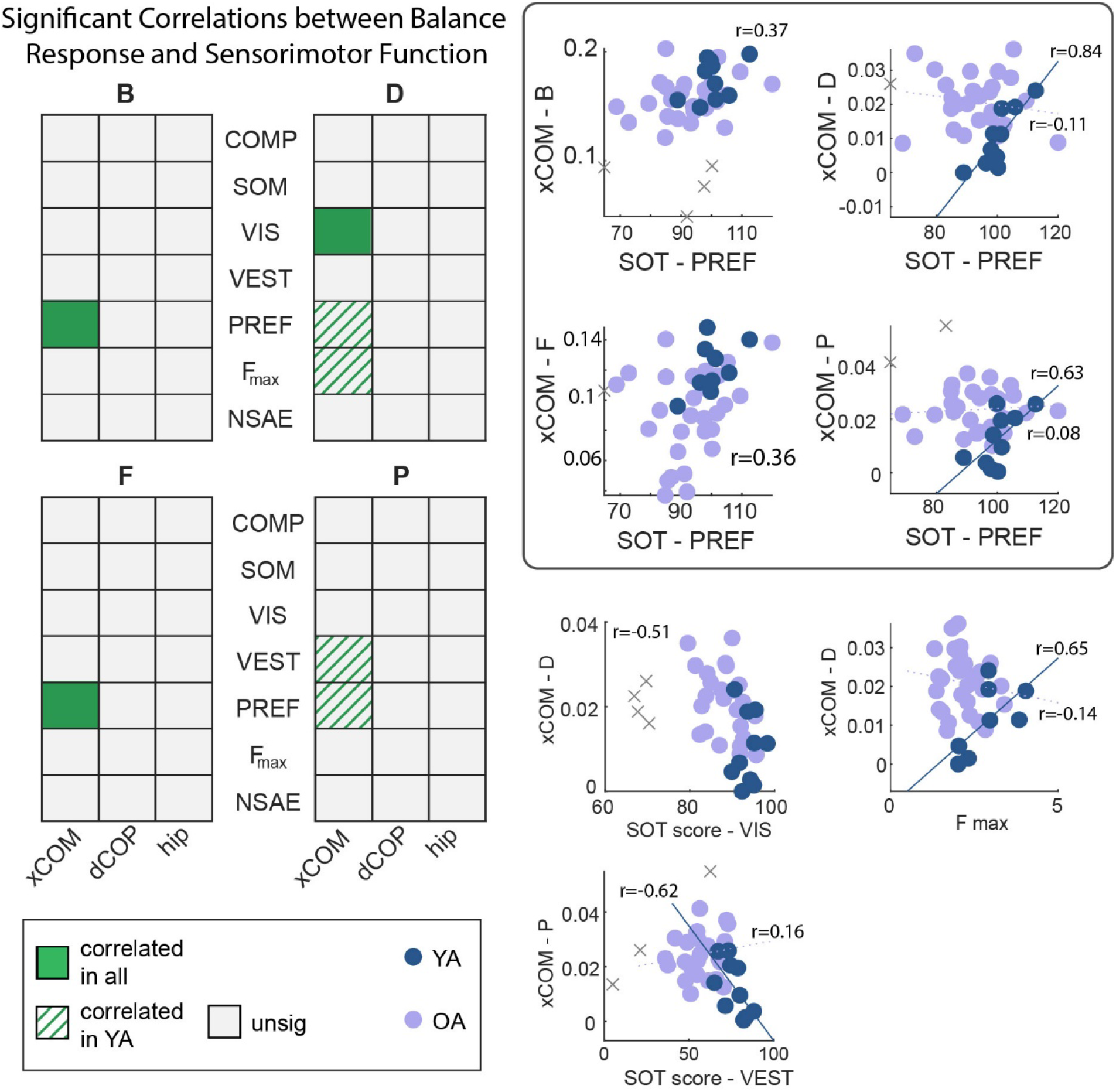
Results of exploratory regression analyses relating measures of sensorimotor function to balance response measures. Out of all comparisons, we identified eight relationships of interest (indicated by green solid fill, hatching, or dots, center). Four of these demonstrated a correlation between balance response and sensory function regardless of age (solid green squares), with the remainder representing relationships in young adults (green hatching) or older adults only (green dot fill). Scatter plots illustrate the comparisons made in each colored square. Here, dark purple indicates young adults and light purple represents older adults. Outliers not included in the regression are marked with ‘x’. We report a single Pearson’s correlation coefficient for correlations across both age groups. For the remaining, solid or dashed lines represent the regression line in each age group, along with the correlation coefficient for each group.

During translations, higher PREF scores were correlated with larger stepping thresholds regardless of age (B: r=0.37, p=0.03, F: r=0.36, p=0.03, Fig 6 left). No other meaningful relationships were identified between sensorimotor function and balance response during translational perturbations.

Stepping threshold during rotations was associated with some measures of sensorimotor function, but only in young adults. Larger stepping thresholds were correlated with higher PREF scores in toe-up (overall model: p<0.001, YA: r=0.84, OA: r=-0.11, Fig. 6 right) and toe-down (overall model: p=0.002, YA: r=0.63, OA: r=0.08, Fig. 6 bottom) rotations, and greater strength (overall model: 0.009, r=0.65, OA: r=-0.14, Fig. 6 right). However, higher VEST scores were associated with lower stepping thresholds during toe-down rotations (overall model: p=0.002, YA: r=-0.62, OA: r=0.16, Fig. 6 bottom).

## DISCUSSION

Healthy older adults step more often than young adults in response to support surface perturbations, not because COM disturbances are larger in older than in young adults but because they step at smaller COM disturbances. In other words, older adults have lower stepping thresholds. Though older adults had significantly worse strength and Sensory Organization Test scores, only the visual preference score (PREF) was associated with stepping threshold across perturbation directions. Individuals who were less able to disregard unreliable visual information were less able or willing to respond to disturbances without stepping. The lower PREF score might reflect a reduced ability to adjust to changing task priorities or sensory inputs and thus greater cognitive rigidity. Older adults might compensate for this greater cognitive rigidity by using a safer balance control strategy, i.e. by taking a step to increase their base of support well before their COM has reached the limits of its current base of support. Though our results were exploratory, they *provide further evidence that cognitive rigidity may be an important research direction and potential therapeutic target*(45).

Older adults step in response to smaller translational disturbances than young adults despite using similar non-stepping kinematic strategies. Compared to young adults, older adults stepped more often in response to smaller COM disturbances (i.e., stepping threshold) during platform translations, consistent with both our hypothesis and previous literature(9,46). However, we found no difference in the magnitudes of ankle or hip responses between age groups. Older adults might thus use similar non-stepping kinematic strategies to respond to the perturbations or alternatively, they may have relied on kinematic strategies not reflected in the ankle or hip movements, such as arm counterrotations.

Platform rotations triggered reactive steps in older adults despite low COM disturbances. Step incidence was low in both age groups during platform rotations, especially during toe-down rotations. Therefore, the peak xCOM deviation in non-stepping responses does not reflect a “stepping threshold”, but rather a measure of maximal COM disturbance. These COM disturbances had values close to zero in young adults but were significantly larger in older adults, indicating that posture was more affected by rotational perturbations in older adults. This is further reflected in the larger ankle and hip responses and the higher step incidence in older than young adults in response to toe-up rotations.

Although there were many differences in sensorimotor function between young and older adults, only few of them were associated with balance strategy. As such, our exploratory analysis provides a clear focus for future research by generating a small set of hypotheses about the relationship between balance control and sensorimotor function. First, older adults may choose to step at smaller COM disruptions in response to translational perturbations because of reduced sensory reweighting ability (low PREF score), rather than reduced physical capacity or reduced acuity of sensory information. Second, toe-up rotational perturbations might induce larger disturbances when visual information is inaccurate (low VIS score). Associations between sensorimotor function and balance control that were only present in young adults should be interpreted with extreme caution given that we included only ten young adults. Although our assessment of strength was limited to a single muscle group, our findings are in line with the lack of consistent associations between strength and balance strategy in literature notwithstanding the large body of research. It is also important to note that SOT does not provide a direct measure of sensory function. Rather, sensory acuity is induced from the contribution of different sensory systems in controlling postural sway.

The ability to tolerate large COM disturbances without stepping might rely on the ability to effectively filter out unreliable sensory information. We found that higher PREF score, reflecting the ability to reject unreliable visual information, might be associated with higher stepping thresholds in response to platform translations, regardless of age. Due to the exploratory nature of our study with many comparisons, these results should be confirmed in a follow up study. However, the correlation between PREF and stepping threshold was the only relationship observed in all four perturbation directions, although only in young adults for rotational perturbations. Being able to quickly and effectively evaluate quality of sensory inputs and rapidly switch between using or disregarding that information (i.e., sensory reweighting) may be important for maintaining balance in unexpected conditions. Our results suggest that individuals with poor sensory reweighting and therefore uncertain incoming sensory information prefer balance strategies that prioritize safety without requiring precise information.

This result is consistent with previous evidence that older adults step without nearing their limits of stability(46), and raises interesting questions about stepping responses: what is the purpose of stepping if there is little threat of instability? Indeed, there is evidence stepping responses in older adults are associated more with whether a disturbance occurred, regardless of the mechanical necessity(46). This is consistent with our finding that older adults step at smaller COM disturbances. In addition, we found that this may be related to susceptibility to unreliable visual input (i.e., PREF score). Poor sensory reweighting might be a sign of reduced cognitive flexibility, i.e. the ability to rapidly switch between modes of thinking or moving(45,47). Reduced set shifting ability is another sign of reduced cognitive flexibility and has been associated with stiffer responses to platform translations, i.e. with increased antagonist co-contraction and smaller COM displacements(45). Stiffening up could also be considered a general strategy to resist translational perturbations. We did not observe stiffer responses here, possibly because we used a mix of translational and rotational perturbations and stiffening up would only help to withstand translational perturbations. In future work we will more specifically assess how set shifting ability is related to sensory reweighting and balance responses.

## CONCLUSIONS

Healthy older adults step more often and at smaller stepping thresholds than young adults, despite using similar magnitude ankle and hip responses. Though older adults performed worse than young adults in measures of sensory and motor function, stepping thresholds were mainly associated with susceptibility to unreliable visual information. Poor sensory reweighting may contribute to and combine with age-related cognitive rigidity, leading to a higher priority on safer strategies. Older adults may resort to stepping, even if a step is not necessary, rather than rely on potentially inaccurate sensory signals to inform a corrective response. Future work should further investigate the relationship between sensory reweighting, cognitive rigidity, and balance.

## GLOSSARY

AP: Anteroposterior
B: Backwards platform translation
COM: Center of mass
COMP: Composite score Sensory Organization Test
COP: Center of pressure
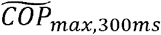: Maximum value of center of pressure in first 300 ms
D: Dorsiflexion (toe-up) platform rotations
F: Forwards platform translations
FES-I: Falls Efficacy Scale-International
Fmax: Maximum knee extension torque
GT: Generated torque
Lmax: Height of center of mass at baseline
NSAE: Normalized standard deviation of absolute error (of torque in torque-matching task)
OA: Older adults
P: Plantarflexion (toe-down) platform rotations
PREF: Visual preference score, Sensory Organization Test
SD: Standard deviation
SOT: Sensory Organization Test
SOM: Somatosensory score, Sensory Organization Test
xCOM: Extrapolated center of mass
*xCOM*_*max,non-stepping*_: Maximum values of extrapolated center of mass in non-stepping trials
VEST: Vestibular score, Sensory Organization Test
VIS: Visual score, Sensory Organization Test
YA: Young adults
*θ*_*AP*_: Estimated anteroposterior postural sway in Sensory Organization Test
*θ*_*AP,max*_: Maximum value of trunk lean
*θ*_*AP,min*_: Minimum value of trunk lean
*θ*_*trunk,max*_: Maximum forward or backward trunk lean during platform perturbations

## ACKNOWLEDGEMENTS

We would like to thank Tom Van Wouwe for the data collection and contributions to design and analysis. We would also like to thank all experiment participants.

## GRANTS

This work was funded by a Research foundation Flanders (FWO) research project (G088420N) to FDG.

## DISCLOSURES

The authors declare that we have no conflicts of interest.

## AUTHOR CONTRIBUTIONS

Hannah D. Carey: analyzed data, interpreted results of experiments, prepared figures, drafted manuscript, edited and revised manuscript

Friedl De Groote: Conceived and designed research, interpreted results of experiments, drafted manuscript, edited and revised manuscript

## Notes

### Competing Interest Statement

The authors have declared no competing interest.

