## Supplementary Material for "Smaller stepping thresholds in older adults might be related to reduced ability to suppress conflicting sensory information"

### Stepping incidence in each perturbation direction

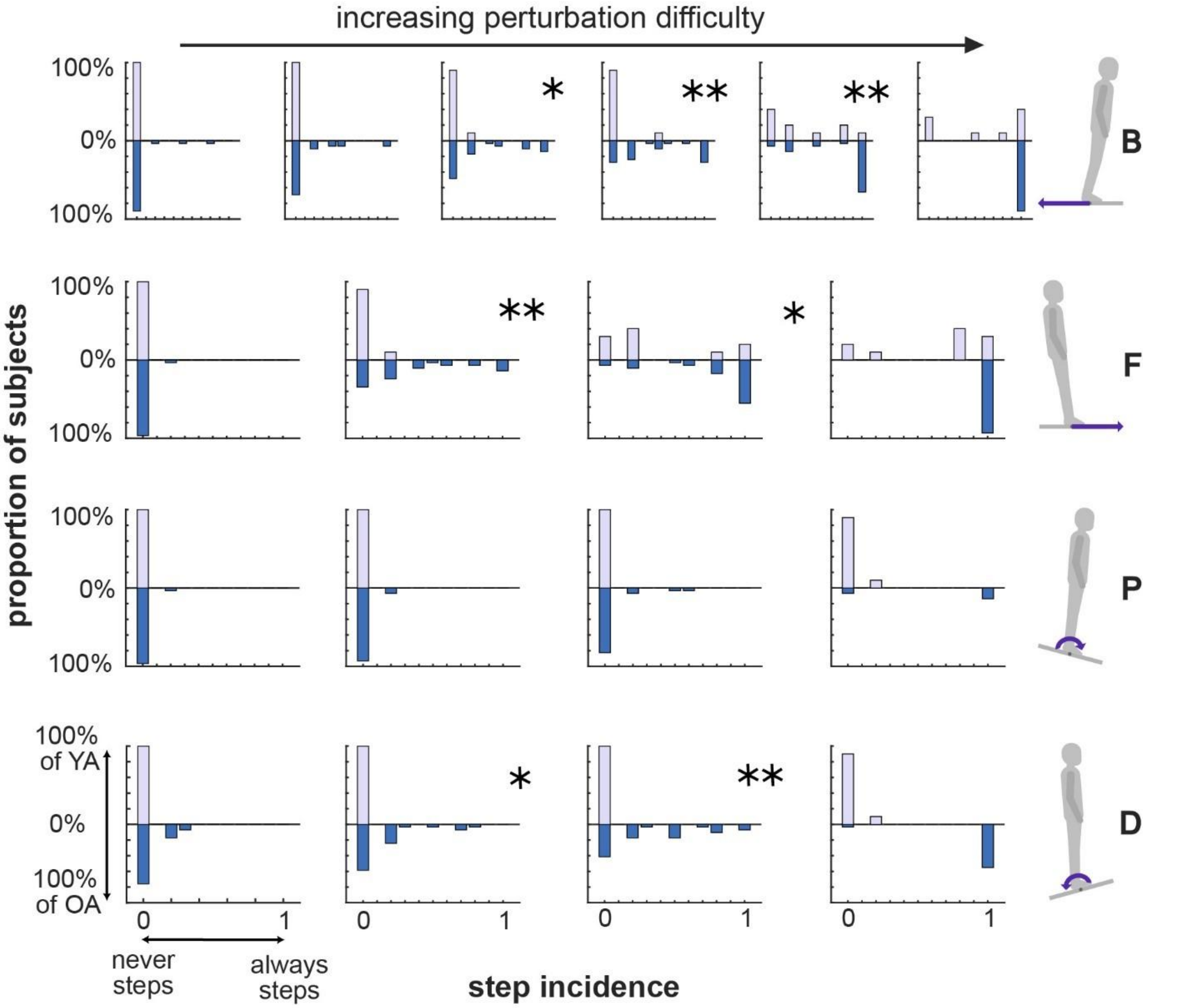


Figure S1. Histograms of step incidence in each perturbation type for young adults (light purple) and older adults (dark blue), including backwards translations (top row), forwards translations (2nd row), toe-down rotations (3rd row), and toe-up rotations (last row). For each plot, bar height indicates the proportion of subjects with a given step incidence. For example, the bottom left plot shows that no young adults stepped during the smallest toe-down rotation, whereas a few older adults sometimes stepped. Stars and p-values indicate a significant difference in step incidence between young and older adults. The most difficult level for each direction was not tested, as many older adults could not complete it.

### S2: Stepping incidence – ANOVA results and post hoc tests

**Repeated Measures ANOVA**

Within Subjects Effects

| | Sum of Squares | df | Mean Square | F | p |
|---|---|---|---|---|---|
| **stepping condition** | 1459 | 3 | 486.5 | 15.96 | <.001 |
| **stepping condition ✻ ageGroup** | 371 | 3 | 123.7 | 4.06 | 0.009 |
| **Residual** | 3200 | 105 | 30.5 | | |

Note. Type 3 Sums of Squares

Between Subjects Effects

| | Sum of Squares | df | Mean Square | F | p |
|---|---|---|---|---|---|
| **ageGroup** | 2.15 | 1 | 2.15 | 0.00576 | 0.940 |
| **Residual** | 13070.19 | 35 | 373.43 | | |

Note. Type 3 Sums of Squares

**Post Hoc Tests**

Post Hoc Comparisons - stepping condition

| Comparison | | | | | | | |
|---|---|---|---|---|---|---|---|
| **stepping condition** | | **stepping condition** | **Mean Difference** | **SE** | **df** | **t** | $p_{tukey}$ |
| **control** | - | **accuracy** | -0.0234 | 0.963 | 35.0 | -0.0243 | 1.000 |

Post Hoc Comparisons - stepping condition

| Comparison | | | | | | | |
|---|---|---|---|---|---|---|---|
| **stepping condition** | | **stepping condition** | **Mean Difference** | **SE** | **df** | **t** | **$p_{tukey}$** |
| | **-** | **speed** | 7.5973 | 1.607 | 35.0 | 4.7268 | <.001 |
| | **-** | **stability** | 0.4188 | 0.824 | 35.0 | 0.5086 | 0.957 |
| **accuracy** | **-** | **speed** | 7.6207 | 1.736 | 35.0 | 4.3895 | <.001 |
| | **-** | **stability** | 0.4422 | 0.822 | 35.0 | 0.5382 | 0.949 |
| **speed** | **-** | **stability** | -7.1785 | 1.621 | 35.0 | -4.4279 | <.001 |

Post Hoc Comparisons - stepping condition ✻ age group

| Comparison | | | | | | | | | |
|---|---|---|---|---|---|---|---|---|---|
| **stepping condition** | **Age group** | | **stepping condition** | **Age group** | **Mean Difference** | **SE** | **df** | **t** | **$p_{tukey}$** |
| **control** | **YA** | - | **control** | **OA** | 2.8260 | 3.47 | 35.0 | 0.8140 | 0.991 |
| | | - | **accuracy** | **YA** | 0.6762 | 1.18 | 35.0 | 0.5707 | 0.999 |
| | | - | **accuracy** | **OA** | 2.1029 | 3.72 | 35.0 | 0.5649 | 0.999 |
| | | - | **speed** | **YA** | 11.8673 | 1.98 | 35.0 | 6.0015 | <.001 |
| | | - | **speed** | **OA** | 6.1533 | 3.62 | 35.0 | 1.7003 | 0.687 |
| | | - | **stability** | **YA** | 1.5981 | 1.01 | 35.0 | 1.5774 | 0.760 |
| | | - | **stability** | **OA** | 2.0655 | 3.53 | 35.0 | 0.5856 | 0.999 |
| | **OA** | - | **accuracy** | **YA** | -2.1497 | 3.63 | 35.0 | -0.5928 | 0.999 |
| | | - | **accuracy** | **OA** | -0.7231 | 1.52 | 35.0 | -0.4761 | 1.000 |
| | | - | **speed** | **YA** | 9.0414 | 3.56 | 35.0 | 2.5383 | 0.213 |

Post Hoc Comparisons - stepping condition ✻ age group

| Comparison | | | | | | | | | |
|---|---|---|---|---|---|---|---|---|---|
| **stepping condition** | **Age group** | | **stepping condition** | **Age group** | **Mean Difference** | **SE** | **df** | **t** | $p_{tukey}$ |
| | | - | **speed** | **OA** | 3.3273 | 2.53 | 35.0 | 1.3128 | 0.888 |
| | | - | **stability** | **YA** | -1.2279 | 3.51 | 35.0 | -0.3503 | 1.000 |
| | | - | **stability** | **OA** | -0.7605 | 1.30 | 35.0 | -0.5857 | 0.999 |
| **accuracy** | **YA** | - | **accuracy** | **OA** | 1.4267 | 3.87 | 35.0 | 0.3689 | 1.000 |
| | | - | **speed** | **YA** | 11.1911 | 2.14 | 35.0 | 5.2396 | <.001 |
| | | - | **speed** | **OA** | 5.4770 | 3.77 | 35.0 | 1.4537 | 0.826 |
| | | - | **stability** | **YA** | 0.9219 | 1.01 | 35.0 | 0.9119 | 0.983 |
| | | - | **stability** | **OA** | 1.3892 | 3.68 | 35.0 | 0.3776 | 1.000 |
| | **OA** | - | **speed** | **YA** | 9.7644 | 3.81 | 35.0 | 2.5648 | 0.203 |
| | | - | **speed** | **OA** | 4.0504 | 2.74 | 35.0 | 1.4795 | 0.813 |
| | | - | **stability** | **YA** | -0.5048 | 3.75 | 35.0 | -0.1345 | 1.000 |
| | | - | **stability** | **OA** | -0.0374 | 1.30 | 35.0 | -0.0289 | 1.000 |
| **speed** | **YA** | - | **speed** | **OA** | -5.7141 | 3.71 | 35.0 | -1.5420 | 0.780 |
| | | - | **stability** | **YA** | -10.2692 | 1.99 | 35.0 | -5.1488 | <.001 |
| | | - | **stability** | **OA** | -9.8019 | 3.62 | 35.0 | -2.7107 | 0.153 |
| | **OA** | - | **stability** | **YA** | -4.5552 | 3.65 | 35.0 | -1.2475 | 0.911 |
| | | - | **stability** | **OA** | -4.0878 | 2.56 | 35.0 | -1.5990 | 0.748 |
| **stability** | **YA** | - | **stability** | **OA** | 0.4674 | 3.56 | 35.0 | 0.1313 | 1.000 |
